# sKL/mKL Transcript Ratio and Protein Localization Define a Species- and Region-Specific Klotho Signature in the CNS and AD Progression

**DOI:** 10.1101/2025.05.13.653838

**Authors:** Rebeca Blanch, Jon Esandi, Ruben Guerrero-Yagüe, Javier del Rey, Joan Roig-Soriano, Alberto Lleo, Eva Carro, Nadine Mestre-Francés, Gina Devau, Isidre Ferrer, Anna Meseguer, Jose Luis Lanciego, Beatriz Almolda, Assumpció Bosch, Miguel Chillón

## Abstract

α-Klotho is a multifunctional protein widely recognized for its anti-aging and neuroprotective properties. This study investigates the expression and localization of the secreted Klotho (s-KL) isoform in the human brain and its potential role in Alzheimer’s disease. Using RT-qPCR, we observed that the s-KL transcript predominates over the membrane-bound KL (m-KL) in multiple brain regions, a pattern consistent in macaques and lemurs. Immunohistochemistry and immunoprecipitation assays confirmed the presence of the s-KL protein in human and mouse brain parenchyma, revealing species-specific cellular localization. In human cerebrospinal fluid (CSF), s-KL constitutes ∼28% of total KL, with levels significantly reduced in mild dementia-AD patients. These findings underscore s-KL’s potential neuroprotective role and highlight its differential regulation and expression during AD progression.

## INTRODUCTION

α-Klotho (KL) is a multifunctional protein widely acknowledged for its anti-aging and neuroprotective properties (Kuro-o et al. 1997; Abraham et al. 2016). Elevated KL levels are associated with improved cognitive function, enhanced healthspan, increased longevity, and greater resistance to oxidative stress across both murine models and humans (Arking et al. 2002; Kurosu et al. 2005; Dubal et al. 2015; Masso et al. 2018). In contrast, decreased KL levels have been observed in the brains of aged mice, rats, and rhesus monkeys, as well as in the cerebrospinal fluid (CSF) of elderly humans compared to young adults (Duce et al. 2008; Semba et al. 2014; Masso et al. 2015). Transgenic KL-deficient mice exhibit an aging phenotype characterized by impaired cognition, motoneuron degeneration, arteriosclerosis, sarcopenia, and premature mortality (Kuro-o et al. 1997). Furthermore, KL-deficient mice exhibit age-related central nervous system (CNS) impairments, such as neuronal degeneration, hypomyelination, disrupted axonal transport, heightened oxidative stress, and reduced levels of dopaminergic neurons, synapses, and synaptic proteins compared to wild-type controls (Uchida et al. 2001; Shiozaki et al. 2008; Kosakai et al. 2011; Chen et al. 2013).

The KL gene (kl) is expressed in a tissue-specific manner, with highest expression levels in the kidney and choroid plexus of the brain. Lower kl expression is observed in skeletal muscle, brain parenchyma, urinary bladder, aorta, colon, small intestine, and various endocrine organs (Kuro-o et al. 1997; Matsumura et al. 1998; Shiraki-Iida et al. 1998). KL is thought to function in a cell-non-autonomous manner, acting as a hormone throughout the organism. It exerts its neuroprotective properties by mitigating oxidative stress, enhancing synaptic plasticity, reducing neuroinflammation, and promoting beneficial autophagy.

The kl gene comprises five exons and expresses two transcript variants. The full-length transcript encodes the membrane-bound form of Klotho (m-KL), which contains a transmembrane domain and two extracellular domains (KL1 and KL2). Proteolytic cleavage of m-KL by specific proteases generates processed KL variants (p-KL) (Chen et al. 2007; Bloch et al. 2009). Alternatively, a second transcript generated via alternative splicing encodes the secreted form of Klotho (s-KL), which consists of the KL1 domain and a unique peptide sequence not present in the other KL isoforms (Matsumura et al. 1998; Shiraki-Iida et al. 1998).

The p-KL and s-KL proteins are collectively referred to as soluble KL, as both are released into biological fluids, including serum, CSF, and urine (Imura et al. 2004; Hu et al. 2011). However, the relative contributions of each variant to the overall soluble KL protein pool remain unclear, partly due to historical limitations in differentiating them. Most existing research has focused on the full-length transcript variants (m-KL and p-KL) or the KL1 domain, with little specificity toward the s-KL isoform. Moreover, the putative receptor for KL has not yet been identified, leaving the precise roles of the s-KL and m-KL isoforms in the CNS unresolved. Nevertheless, some reports suggest a crucial role of s-KL in CNS aging and Alzheimer’s disease (AD) pathology in murine models (Masso et al. 2015; Masso et al. 2018; Roig-Soriano et al. 2022). Its unique properties as a secreted, soluble protein, and its distinct peptidic sequence, may confer unique functionalities. Notably, reduced levels of soluble KL have been observed in the CSF of human AD patients compared to healthy controls (Semba et al. 2014), along with decreased KL expression in various AD mouse models (Dubal et al. 2015; Masso et al. 2015; Kuang et al. 2017). In addition, KL administration in an AD mouse model has been shown to improve cognitive symptoms and AD neuropathology (Zeng et al. 2019). As most studies have primarily focused on the full-length KL variant and its processed forms, this study aimed to characterize the expression of s-KL in the healthy human CNS, and under AD-related pathological conditions, as well to explore its potential association with AD risk factors.

## Results

### Gene expression ratio of s-KL vs m-KL in the human brain

Our analysis revealed that the primary KL transcript variant detected in both human and mouse kidneys was m-KL, with approximately 3-fold difference over s-KL in humans and over 30-fold in mice (Fig. 1 A and B), in accordance with prior research (Matsumura et al. 1998; Mencke et al. 2017; Olauson et al. 2017).

**Figure 1.**
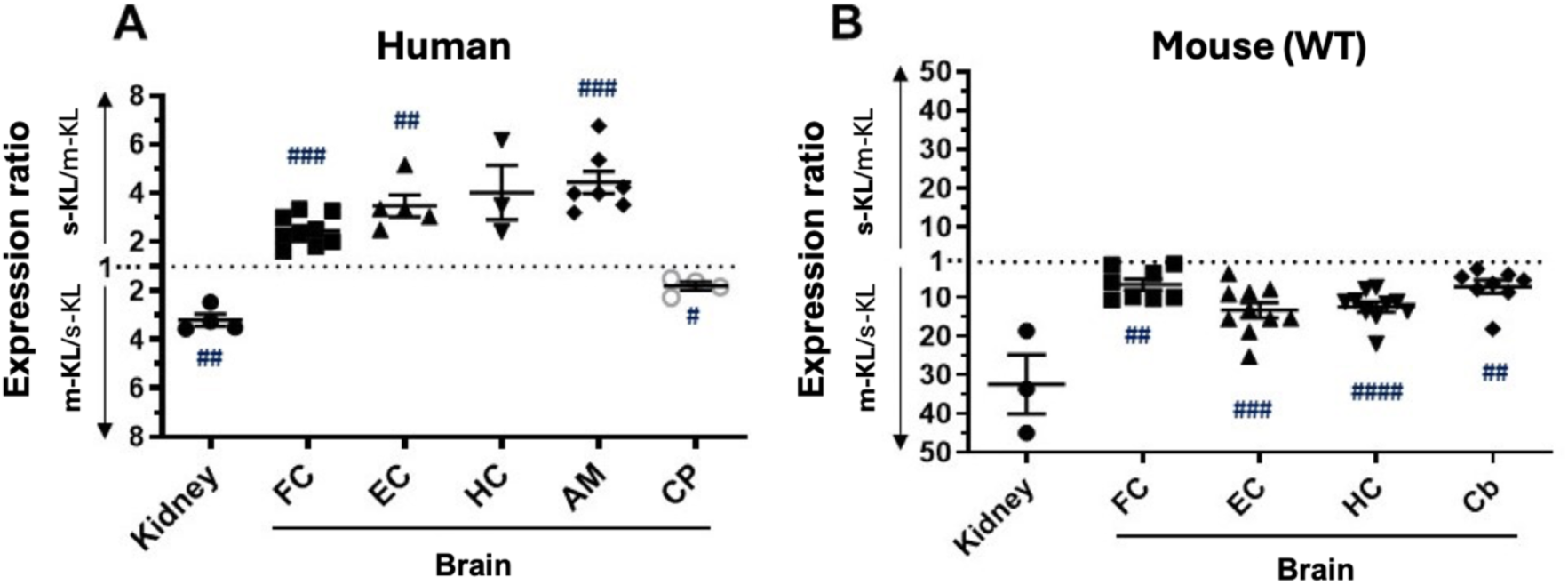
Gene expression ratio of s-KL *vs* m-KL in (A) human kidney and brain (B) mouse kidney and brain. Individual dots represent the ratio for each subject in each area. Mean±SEM plotted per area. The one-sample t-test was used to determine whether a mean ratio was different from 1, in which case the relative proportion of the KL variants would be different (statistical significance indicated with: # p<0.05, ## p<0.01, ### p<0.001, #### p<0.0001). FC: frontal cortex, EC: entorhinal cortex, HC: hippocampus, AM: amygdala. CP: choroid plexus. Cb: cerebellum. Sample size (n): **A**: FC(9), EC(5), HC(3), AM(7), CP(4). **B**: Kidney(3), FC(8), EC(10), HC(10), Cb(8).

In the human brain, our RT-qPCR results revealed, for the first time, the presence of the s-KL transcript in various regions of the CNS parenchyma, including the frontal and entorhinal cortex, hippocampus, amygdala, and choroid plexus (Fig. 1A). Notably, the expression of the s-KL isoform transcript was 2.5 to 4.4 times higher than that of m-KL in all the areas analyzed, reaching statistical significance in the frontal cortex, entorhinal cortex, and amygdala. In contrast, in the choroid plexus, the relative abundance of s-KL compared to m-KL was inverted, with the m-KL transcript being significantly more abundant than s-KL (1.8-fold).

To assess whether this ratio between s-KL and m-KL transcript isoforms was evolutionarily conserved, we analyzed the expression of these isoforms in other species, including mice, the non-human primate lemur (*Microcebus murinus*), and the rhesus monkey. Our analysis revealed that in mice, the m-KL transcript was significantly more abundant than s-KL in all brain regions examined, including the frontal cortex (6.7-fold), entorhinal cortex (13.3-fold), hippocampus (12.5-fold), and cerebellum (7.3-fold) (Fig. 1B). In lemur brain tissue, the s-KL to m-KL ratio was close to 1, with no statistical significance detected (One sample t-test, hipp=1), indicating similar expression levels of s-KL and m-KL in this species (Supp. Fig. 1A). Finally, in the rhesus monkey, similar to humans, the s-KL variant showed greater expression than m-KL by more than 146-fold in all analyzed brain regions: superior frontal gyrus (909-fold), precentral gyrus (412-fold), entorhinal cortex (759-fold), hippocampus (146-fold), and cerebellum (1169-fold). However, no statistical significance difference was detected, potentially due to high variability and the limited sample size (n=3 per brain region) (Supp. Fig. 1B).

### Detection of the s-KL protein in different brain areas across different species

Using immunofluorescence (IF) techniques, we detected the s-KL protein in human brain tissues. We observed s-KL colocalization with total KL in both hippocampus and amygdala, as well as a distinctive punctate intracellular pattern in regions where colocalization patterns did not align (Fig. 2A). Additionally, s-KL protein also colocalizes with total Klotho in different areas of the mouse brain (Supp. Fig. 2A). Interestingly, in mice, s-KL predominantly localizes to neurons, similar to m-KL, whereas in non-human primates the common KL1 domain (there is no specific antibody for detecting lemur’s s-KL) is detected in both neurons and astrocytes (Supp. Fig. 2B). In contrast, in human brain, s-KL was primarily detected in astrocytes, though not exclusively, with minimal to negligible detection in neurons (Fig. 2B). This underscores not only species-specific expression differences but also suggests an evolutionary divergence in cellular expression profiles.

**Figure 2.**
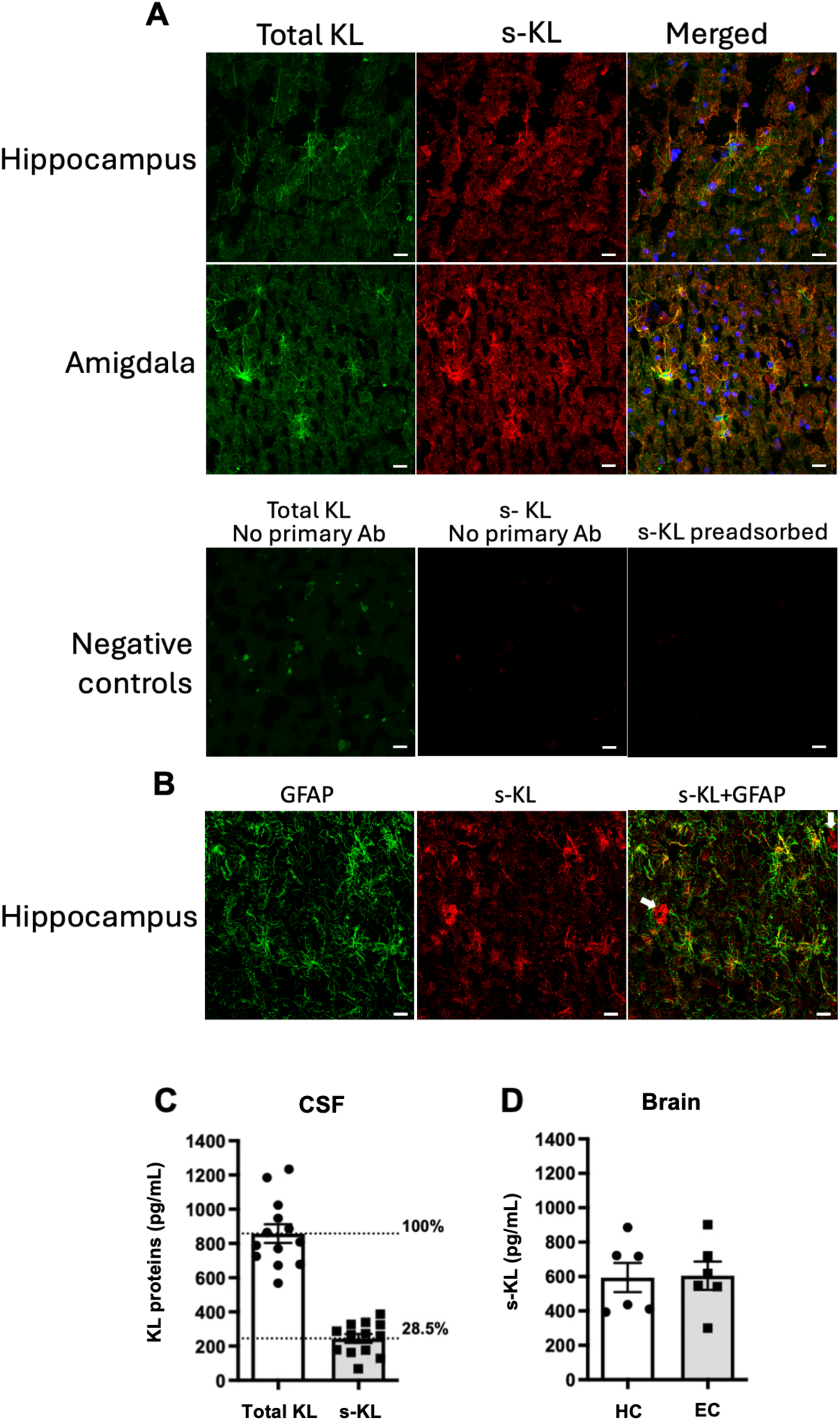
Total klotho and s-KL expression in CSF and different human brain areas of healthy individuals. Representative images of double immunolabelling sections from human hippocampus and amygdala for total klotho and s-KL **(A)**; or for s-KL and GFAP **(B)**. Negative controls were done by incubating sections without the primary antibodies or by incubating with the pre-adsorbed antibody. White arrows in (B) point to the few neurons expressing s-KL, while yellow labelling indicates s-KL positive astrocytes. Bar scale size is 10μm. Quantification of total soluble klotho (KL1) and s-KL in CSF **(C)** and homogenized brain tissue (HC: hippocampus, EC: entorhinal cortex) **(D)** by ELISA.

In mouse brain parenchyma, the presence of the s-KL protein was also confirmed using immunoprecipitation assays combined with Western blotting. Specifically, immunoprecipitation with the BAF1819 and KM2076 antibodies, both targeting the common KL1 domain, followed by detection with the K113 antibody specific for the unique murine s-KL region (Supp. Fig. 2C and 2D). Similarly, reciprocal immunoprecipitation using the K113 antibody, and subsequent WB detection with the BAF1819 and KM2076 antibodies further corroborated the findings (Supp. Fig. 2E and 2F). These results conclusively demonstrate that the s-KL protein is actively produced within the mouse brain parenchyma.

To further characterize KL expression in the healthy human CNS, we quantified the protein levels of KL variants using two commercially available ELISA kits: one detecting the KL1 domain (present in all KL variants, referred to as total KL), and the other specifically detecting the s-KL protein. In human CSF, the s-KL isoform represented approximately 28.5% of the total soluble Klotho pool, indicating that soluble KL proteins in the CSF pool primarily derive from the m-KL variant (Fig. 2C). This agrees with the gene expression levels observed in the choroid plexus -the tissue where the CSF proteins are produced-where m-KL variant expression was higher than that of s-KL (Fig. 1A).

Interestingly, within human brain parenchyma (hippocampus and entorhinal cortex), s-KL protein levels were approximately 3 to 4-fold higher than in CSF (Fig. 2D), aligning with gene expression data indicating a higher expression of s-KL in parenchymal brain regions (Fig. 1A). Unfortunately, we could not determine the relative proportion of s-KL versus total KL in human brain parenchyma due to technical constraints associated with the commercial ELISA kit for total Klotho.

Overall, these findings highlight the distinct expression pattern and complex spatial distribution of the KL variants within the human brain, suggesting a different role for s-KL in neurophysiological processes. Of note, further investigation is needed to elucidate the mechanisms regulating s-KL expression and its functional implications in the human CNS.

### Gene expression ratio of s-KL vs m-KL across stages of Alzheimer’s disease

After confirming the presence of the s-KL transcript and protein in the human brain, we aimed to determine whether the s-KL/m-KL ratio is altered across neuropathological stages of AD. For this purpose, we analyzed by RT-qPCR the relative expression of s-KL versus m-KL in brain regions of AD patients, who were categorized post-mortem according to the Braak staging criteria into: AD I/II (clinically silent AD), AD III/IV (incipient AD), and AD IV-VI (incipient and fully developed AD) (Braak and Braak 1991).

As in healthy individuals, the s-KL isoform was the predominant variant in all parenchymal brain regions analyzed in AD patients (Fig. 3). However, in contrast to healthy controls, the relative abundance of s-KL in the choroid plexus was comparable to that of m-KL transcripts.

**Figure 3.**
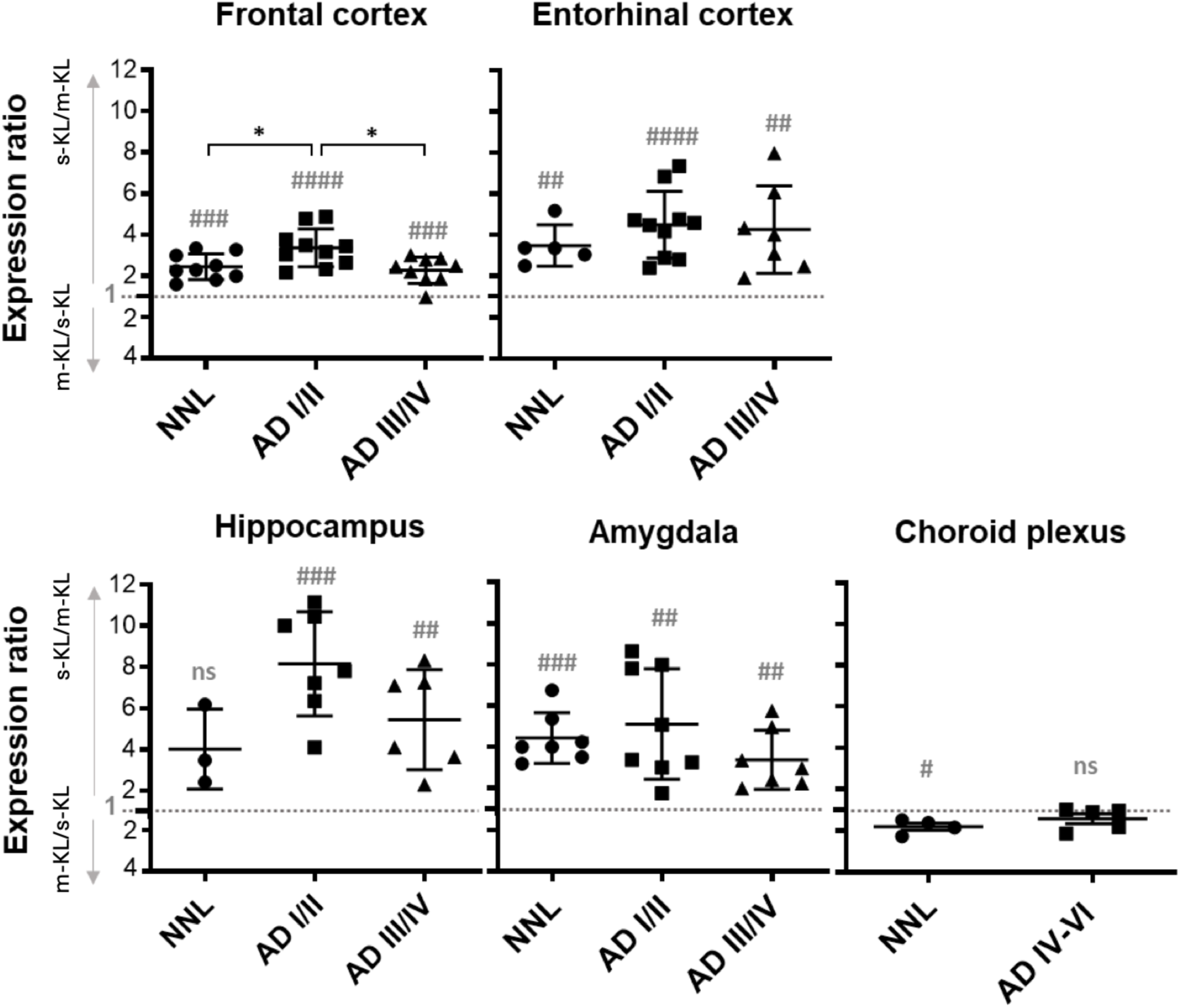
Gene expression ratio of s-KL vs m-KL in brain areas of aged NNL (Braak 0) and AD patients (ADI/II and ADIII/IV) classified according to the Braak staging criteria. Individual dots represent the ratio for each subject, plotted as Mean±SEM. One-sample t-test was used to determine whether a mean ratio was different from 1, in which case the relative proportion of s-KL vs m-KL would be different in a particular area (statistical significance indicated with # p<0.05, ## p<0.01, ### p<0.001). One-way ANOVA and Tukey post-hoc tests were used to detect differences in the mean ratio between groups in each area (statistical significance indicated with *p<0.05). NNL: Donors with no neuropathological lesions. Sample size per area and group (n): FC (NNL:9, ADI/II:10, ADIII/IV:9); EC (NNL:5, ADI/II:10, ADIII/IV:7); HC (NNL:3, ADI/II:7, ADIII/IV:6); AM (NNL:7, ADI/II:8, ADIII/IV:7); CP (NNL:4, ADIV-VI:5).

Notably, the s-KL/m-KL ratio was higher in clinically silent AD patients (AD I/II) compared to the healthy group, particularly in the frontal cortex and hippocampus−regions heavily associated with AD pathology.

To further investigate differences in relative s-KL vs m-KL expression in AD patients, the transcription levels of the two KL isoforms were normalized to two reference genes, cytochrome c1 (CYC1) and ubiquitin conjugating enzyme E2 D2 (UBE2D2), as recommended (Rydbirk et al. 2016) (Supplementary Fig 3). Although no statistically significant differences were observed, some trends could be distinguished. First, s-KL expression was slightly increased in the hippocampus, amygdala, and choroid plexus of AD patients compared to healthy donors, while in the frontal cortex, s-KL tended to decrease. Second, m-KL showed a slight decrease in the frontal and entorhinal cortex of AD patients compared to healthy donors, while substantial variability masked a potential trend toward increased m-KL expression in the amygdala and choroid plexus, particularly in advanced AD stages (AD III/IV and AD IV-VI, respectively).

The elevated s-KL/m-KL expression ratio observed in the frontal cortex of the AD I/II group may be attributed to a decrease in m-KL transcript levels, whereas in the hippocampus, it seemed a result from increased s-KL expression combined with decreased m-KL expression. In the choroid plexus of AD patients, the gene expression of both m-KL and s-KL transcripts was elevated compared to healthy individuals, although the increase in s-KL seemed more pronounced, resulting in similar transcript levels of m-KL and s-KL in AD patients.

### Protein levels of s-KL and total KL in CSF and brain of AD patients

The protein levels of total soluble KL and secreted KL were evaluated in the CSF of cognitively unimpaired (CU) individuals, mild cognitive impairment (MCI), or mild dementia (MD) patients (Table 3). The levels of total KL in CSF were significantly reduced in AD dementia patients (655.2 ± 231.9 pg/mL) compared to CU controls (857.8 ± 197.5 pg/mL), while MCI patients showed similar levels (870.5 ± 246.9 pg/mL) to the CU group (Fig. 4A). Similar to total KL, levels of s-KL in CSF tended to decrease in AD dementia patients (Fig 4H). Next, we studied the correlation of CSF KL levels with relevant AD risk factors such as sex, age and APOE genotype.

**Figure 4.**
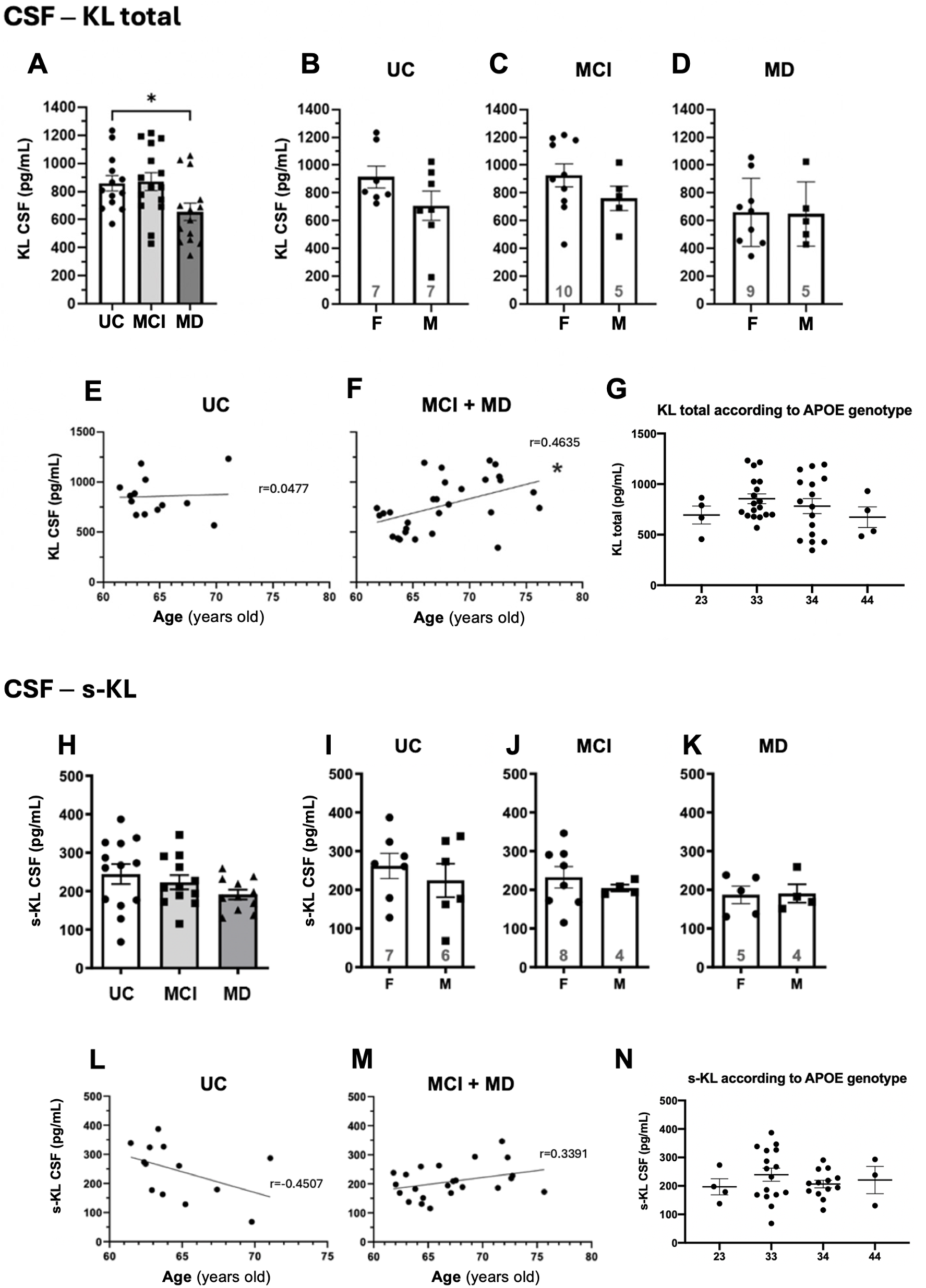
KL protein levels in human CSF. (A-G, total soluble Klotho (p-KL and s-KL); H-N, s-KL). **A, H:** Quantification in aged cognitive unimpaired individuals (CU) and AD patients with mild cognitive impairment (MCI) and mild dementia (MD). One-way ANOVA and Dunnett’s post hoc tests were performed to detect differences between the mean of each AD group and the cognitively healthy controls (*p<0.05). Sample size (n): CU (n=13), MCI (n=15), MD (n=14). **B-D, I-K**: Comparison of KL-CSF levels in women *vs* men of different clinical status. F: women, M: men. Differences between F and M analyzed by the t-test. Mean±SEM represented. Sample size indicated within the graph bars. **E-F, L-M:** Correlation of KL-CSF and s-KL-CSF levels with age (r = Pearson correlation coefficient). Statistically significant linear dependence denoted by *p<0.05 and ***p<0.001. Sample size (n): CU (13); AD (29). **G, N**: KL-CSF levels in all subjects according to the presence of the APOE4 allele without clinical distinction. 23,33,34,44: APOE alleles. Individual dots represent KL levels of each subject. Mean±SEM is represented. Differences between two groups analyzed by t-test (**p<0.01). Sample size indicated within the graph bars.

No significant sex-based differences were observed in total KL or s-KL levels across clinical groups (Fig. 4B-D, 4I-K). Of note, in AD patients s-KL accounted for ∼28% of total soluble KL in CSF, consistent CU individuals.

Age correlation analysis revealed no age-related variation in total KL levels in CU individuals over 61 years (Pearson test, Fig. 4E). However, AD patients showed a significant positive correlation (r=0.4635, p=0.011), indicating elevated total KL with age (Fig. 4F). While s-KL levels in CU individuals displayed a non-significant decline with age (r=-0.4507, p=0.122; Fig. 4L), s-KL levels in AD patients with MCI or mild dementia showed a slight age-related increase (Fig. 4M), similar to total soluble KL.

APOE genotype analysis revealed similar CSF levels for both, total soluble KL protein (2/3 carriers: 694.5 ± 178.5 pg/mL; 3/3 carriers: 855.2 ± 204.8 pg/mL; 3/4 carriers 782,5 ± 297.7 pg/mL; and 4/4 carriers 672.7 ± 204.3 pg/mL; Fig4G) and s-KL protein (2/3 carriers: 197.0 ± 56.6 pg/mL; 3/3 carriers: 239.3 ± 91.5 pg/mL; 3/4 carriers 206.5 ± 47.6 pg/mL; and 4/4 carriers 220.6 ± 82.7 pg/mL; Fig4N). Also, s-KL versus total KL ratio was constant (around 28%) regardless the APOE allele. Stratification by clinical groups showed similar trends, but statistical significance was limited by sample size.

Finally, we assessed the levels of the s-KL protein in the hippocampus and entorhinal cortex, two brain regions heavily affected by AD pathology. Elevated s-KL protein levels were observed in clinically asymptomatic AD patients (AD I/II) compared to healthy individuals, with statistical significance observed in the entorhinal cortex (Fig. 5). This observation aligns with the results of gene expression analysis in brain regions, which showed increased s-KL/m-KL ratio in clinically asymptomatic AD patients compared to healthy individuals, both in the hippocampus and entorhinal cortex.

**Figure 5.**
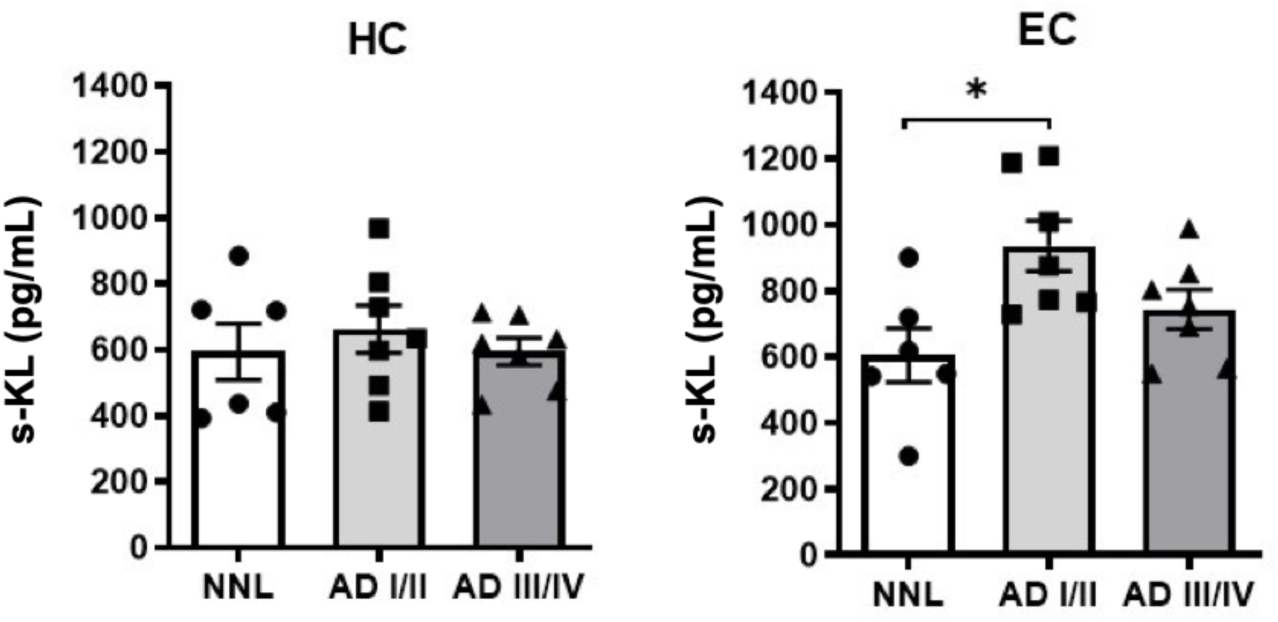
s-KL protein levels in human Hippocampus and Enthorinal Cortex. ELISA of s-KL from Alzheimer’s patients and healthy controls (NNL). One-way ANOVA and Tukey post-hoc tests were used to detect differences in the mean ratio between groups (statistical significance indicated with *p<0.05). Sample size (n): (NNL:6, ADI/II:7, ADIII/IV:7).

## Discussion

In this study, we characterized, for the first time, the relative gene expression of s-KL and m-KL transcripts across various regions of the human brain. Notably, s-KL displayed higher expression levels than m-KL in the frontal cortex, entorhinal cortex, hippocampus, and amygdala, areas involved in cognition, learning, memory, and anxiety. In contrast, the choroid plexus presented an inverted s-KL to m-KL ratio compared to other brain regions, indicating region-specific expression dynamics.

Further comparative analysis across other mammalian species, including mice, lemurs, and rhesus monkeys, revealed diverse expression patterns of s-KL and m-KL transcripts in the brain. In mice, m-KL transcript predominated in all brain regions examined, consistent with previous findings (Shiraki-Iida et al. 1998; Bektas et al. 2004; Masso et al. 2015). In lemur’s brain, the expression ratio of s-KL to m-KL was approximately 1, suggesting similar expression levels of both transcript variants. In contrast, rhesus monkeys showed a similar pattern to humans, with significantly higher expression levels of s-KL compared to m-KL across various brain regions, despite high variability potentially due to the limited sample size. The inverted ratio of s-KL to m-KL expression in the brains of mice and humans could have relevant implications in terms of translatability, considering the widespread use of mouse models in preclinical neurology research. These findings emphasize the relevance of s-KL in the brain parenchyma of both human and non-human primates, while in kidney, m-KL seems to play a more relevant role, being the highest expressed variant across species.

Prior to this work, the s-KL transcript had been detected in human kidney and hippocampus, where the *kl* gene was first identified by Matsumura et al. (Matsumura et al. 1998). They documented the prevalence of the s-KL transcript variant over m-KL in various tissues, including the kidney, a finding that diverges from recent studies (Mencke et al. 2017) and our own observations. Matsumura et al. employed an RNAse protection assay to compare s-KL and m-KL expression in clones isolated from cDNA human libraries, examining 6 kidney clones and 4 hippocampal clones, whereas our study employed RT-qPCR analysis of over 100 mRNA samples from human tissues, thus providing more accurate results. Notably, existing RNA-Seq databases lack the capability to distinguish between the expression of full-length m-KL transcripts and alternative s-KL transcripts (Olauson et al. 2017).

Collectively, our findings, along with previous data (Matsumura et al. 1998; Ohyama et al. 1998; Shiraki-Iida et al. 1998; Clinton et al. 2013), suggest that the expression pattern of s-KL and m-KL is heterogeneous across different tissues and species, which could involve potential differential regulation of KL gene expression and/or distinct functions of s-KL and m-KL across species and organs.

A battery of immunoprecipitation, IHC and ELISA analysis further demonstrated the differential expression and localization of s-KL within the human brain. Notably, our findings reveal the presence of the s-KL protein not only within the human brain parenchyma, where the s-KL transcript predominates, but also in the mouse brain, where m-KL is the predominant isoform. This observation is particularly significant, as the only previous study investigating s-KL expression in human cells reported that the alternative s-KL transcript a target for non-sense mediated decay (NMD) and did not translate into protein in the kidney (Mencke et al. 2017). It is noteworthy that Mencke et al. conducted their investigations primarily in in vitro cellular and tissue kidney models. Therefore, while their hypothesis may be relevant in those specific contexts, we hypothesize that s-KL as a protein plays a more relevant role in the human central nervous system (CNS). While further investigation is needed to clarify s-KL translation and secretion in the kidney, our research demonstrates that in the brain, s-KL is not only highly expressed, but is also actively translated into the s-KL protein.

Additionally, the observation of s-KL protein colocalization with total Klotho in different brain areas may indicate potential interaction between the isoforms. Interestingly, species-specific differences are noted in the cellular localization of s-KL, with predominant expression in astrocytes in humans, neurons in mice, and both neurons and astrocytes in non-human primates. Such differences not only highlight the complexity of Klotho biology but also point out divergent evolutionary trajectories in cellular expression profiles, which may be linked to species-specific neurophysiological adaptations.

Nonetheless, this study also reveals limitations and areas for future investigation. Technical limitations of current ELISA kits prevent direct determination of the relative proportion of s-KL to total KL in brain parenchyma, highlighting the need for improved methodologies to address this gap. Furthermore, the functional implications of region-specific s-KL expression in the brain remain largely unexplored. Given the potential role of KL in various neurophysiological processes such as neuroprotection, synaptic plasticity, and cognition, uncovering the specific functions and regulatory mechanisms of s-KL in the brain is a critical direction for future research.

In AD patients, s-KL was the predominant transcript across all examined parenchymal brain regions—including the frontal cortex, entorhinal cortex, hippocampus, and amygdala—mirroring the expression patterns observed in healthy individuals. Notably, clinically silent AD patients (AD I/II) had an elevated s-KL/m-KL ratio in the frontal cortex and hippocampus compared to healthy controls. This observation is particularly relevant as these brain regions are affected in the initial stages of AD pathology, which involve amyloid-β (Aβ) plaque formation in the isocortex and neurofibrillary tangle (NFT) deposition in the allocortex of the medial temporal lobe, including the hippocampus (Serrano-Pozo et al. 2011). In the choroid plexus of AD patients, the expression levels of s-KL and m-KL were similar, in contrast to healthy individuals where m-KL predominated.

The observed alterations in the s-KL/m-KL ratio in AD patients may reflect a compensatory mechanism to elevate s-KL levels in response to AD pathology, thereby suggesting a putative neuroprotective role for s-KL. It is worth mentioning that the choroid plexus donors were at more advanced AD stages (IV-VI) compared to those of the parenchymal areas (I-IV), thus further analysis is needed using samples from all AD stages to better characterize KL expression dynamics during AD progression.

In an AD-like lemur model exhibiting pathology reminiscent of AD, both s-KL and m-KL transcripts had approximately a 3-fold decrease in the brain compared to healthy subjects (G. Devau, and N. Mestres-Francés, personal communication). Although this variation was more pronounced in lemurs than in human patients, the relative expression of s-KL versus m-KL remained similar in the cerebral cortex of this AD-like lemur model.

The analysis of total soluble KL protein levels in CSF revealed a significant decrease in AD dementia patients compared to CU individuals, whereas MCI-AD patients had total soluble KL CSF levels similar to the CU controls. These findings align with previous reports (Semba et al. 2014; Grontvedt et al. 2022), although those studies did not stratify AD patients based on clinical stage.

KL is well known as an anti-aging protein (Kim et al. 2015) and in the absence of neuropathology, levels of total soluble KL in CSF were previously reported to decrease with advancing age (Semba et al. 2014; Grontvedt et al. 2022). However, in agreement with others(Kundu et al. 2022), we did not observe this correlation in the group of CU individuals (Fig. 4E). Interestingly, we did observe that s-KL CSF levels tended to be decreased with age in CU individuals (Fig. 4L). Therefore, the conflicting results regarding KL-CSF and age correlation could be explained by the fact that the decrease in the soluble KL pool is due to a reduction in the s-KL variant, whereas the previous studies have not specifically analyzed this isoform but the total soluble KL pool.

For individuals with early-stage AD, a positive correlation was observed between total KL CSF levels and age; however, the analysis of s-KL revealed a positive but non-significant trend, requiring caution due to the limited sample size. Correspondingly, in the choroid plexus of AD patients, we detected increased kl gene expression. Another study involving adult patients (aged 41 to 81 years) revealed a significant age-related decrease in total soluble KL-CSF levels in the control group, but not in the AD-MCI or AD-Dementia groups (Grontvedt et al. 2022). Hence, it appears that the positive correlation of KL CSF with age in AD patients may be only associated to elderly adults at early stages of the disease, suggesting the need for future studies stratifying samples according to age groups (adults <61 vs elderly >61) and AD-stages to better comprehend this association.

Overall, these observations could be attributed to a combination of two phenomena. Firstly, age-dependent decline of KL seems accelerated in the presence of AD pathology, resulting in lower total soluble KL CSF mean levels in dementia AD patients compared to CU age-matched individuals. Prior research in rhesus monkey reported a diminished KL expression as a consequence of the age-dependent methylation and silencing of the KL promoter (King et al. 2012). Given that epigenetic mechanisms are altered in the presence of AD pathology (Bellver-Sanchis et al. 2021; Yang et al. 2023), KL downregulation may occur prematurely, as suggested by some authors(Grontvedt et al. 2022). However, whether KL dysregulation contributes to AD onset or is a consequence of the pathology remains unclear. Secondly, we hypothesize that KL upregulation in the elderly population with mild AD may serve as a neuroprotective mechanism to counteract AD pathology, suggesting a compensatory mechanism to mitigate age-dependent AD progression and/or delay pathology onset. A comparable upregulation mechanism at early AD stages has been suggested for other neuroprotective proteins, such as TREM2, which enhances Aβ phagocytosis by microglia (Kim et al. 2017).

Despite the interaction between KL and APOE proteins remains elusive, it is plausible that increased KL levels could mitigate the deleterious effects of the APOE4 allele, a strong genetic risk factor for late-onset AD (Verghese et al. 2011), thereby protecting against AD development and/or progression. Thus, one study in AD-dementia patients reported lower KL-CSF levels in APOE4 carriers compared to noncarriers, although only a small sample size (n=4) for the non-APOE4 group (Grontvedt et al. 2022) was analyzed. In contrast, we observed no correlation between the APOE4 allele and the total soluble KL and s-KL levels in CSF, independent of clinical status, which agrees with different studies in serum and CSF (Kundu et al. 2022; Shibata et al. 2025).

Regarding sex, no significant differences in total soluble KL CSF levels were detected between men and women of advanced age in the entire sample or in groups stratified according to clinical stage, consistent with previous works (Grontvedt et al. 2022; Kundu et al. 2022). However, conflicting results have been reported in other studies analyzing total soluble KL CSF levels according to sex: in a risk-enriched cohort for AD, total soluble KL levels were higher in females compared to males (Gaitan et al. 2022), whereas Semba et al. reported higher total soluble KL CSF levels in males (Semba et al. 2014). Notably, none of these studies stratified the data according to AD status and age, unlike our analysis conducted in elderly individuals (older than 61 years) and stratified according to AD status. An important limitation of our study was the relatively low sample size, especially in the stratified analysis, and therefore, further analyses with larger sample sizes considering AD status, age, and other AD risk factors are needed.

In conclusion, this study is the first to detect and quantify both the s-KL transcript variant and the s-KL protein within the human CNS. Our findings reveal an intricate spatial distribution and distinct expression patterns of s-KL across different brain regions, suggesting a potential role for s-KL in neurophysiological processes. The observed regional and species-specific variations in s-KL and m-KL expression and protein levels emphasize the need for further research to elucidate the functional implications and regulatory mechanisms that govern the expression of the KL variants in the CNS, particularly in the context of AD and other neurological disorders.

### Material and methods

### Human samples

Human biological samples were obtained from different sources in accordance with the guidelines of Spanish legislation (*Real Decreto de Biobancos* 1716/2011).

#### Brain

Post-mortem brain specimens from human donors were provided by Dr. Isidre Ferrer from the Hospital de Bellvitge Biobanc, located in Barcelona, Spain. These specimens comprised four distinct brain regions: the frontal cortex (FC), entorhinal cortex (EC), hippocampus (HC), and amygdala (AM). Patients diagnosed with Alzheimer’s disease (AD) were identified during brain autopsy, with varying severity stages of the disease classified according to the Braak staging criteria (Braak and Braak 1991), and subsequently divided into mild (AD I/II), and moderate (AD III/IV) groups. Only individuals lacking comorbidities were included in the study. Donors with no neuropathological lesions (NNL) were utilized as controls. The distribution of samples analyzed per group is presented in Table 1. Brain specimens were preserved at −80°C. Tissue fragments weighing approximately 100 mg were dissected at 4°C, with careful avoidance of white matter, blood vessels, and hematomas, to facilitate RNA and protein isolation.

**Table 1.** *Post-mortem* human brain samples.

| Diagnosis | Average age (y/o) | Gender (n) |  | Brain areas available (n) |  |  |  |
| --- | --- | --- | --- | --- | --- | --- | --- |
|  |  | F | M | FC | EC | HC | AM |
| NNL | 48.4 | 4 | 6 | 10 | 9 | 6 | 8 |
| AD I/II | 70.0 | 3 | 7 | 10 | 10 | 10 | 10 |
| AD III/IV | 78.4 | 2 | 8 | 9 | 10 | 8 | 9 |

Diagnosis according to Braak criteria. F: females, M: males. y/o: years old. n: number of samples.

#### Kidney

RNA samples from human kidney were provided by Dr Anna Meseguer *(Vall d’Hebron Research Institute,* Barcelona, Spain). The analyzed donors had not been diagnosed with any pathology (n=14).

#### Choroid plexus

RNA samples of human choroid plexus were provided by Dr. Eva Carro (12 de Octubre Hospital Research Institute, Madrid, Spain). These samples were obtained from AD patients staged IV-VI (according to Braak criteria) and non-neuropathological lesions (NNL) individuals (see Table 2).

**Table 2.** Choroid plexus samples from human donors.

| Diagnosis | Average age (y/o) | Gender (n) |  | Total (n) |
| --- | --- | --- | --- | --- |
|  |  | F | M |  |
| NNL | 76.2 | 2 | 3 | 5 |
| AD IV-VI | 75.8 | 2 | 3 | 5 |

Diagnosis according to Braak criteria. F: women, M: males. y/o: years old. n: number of samples.

**Table 3.**
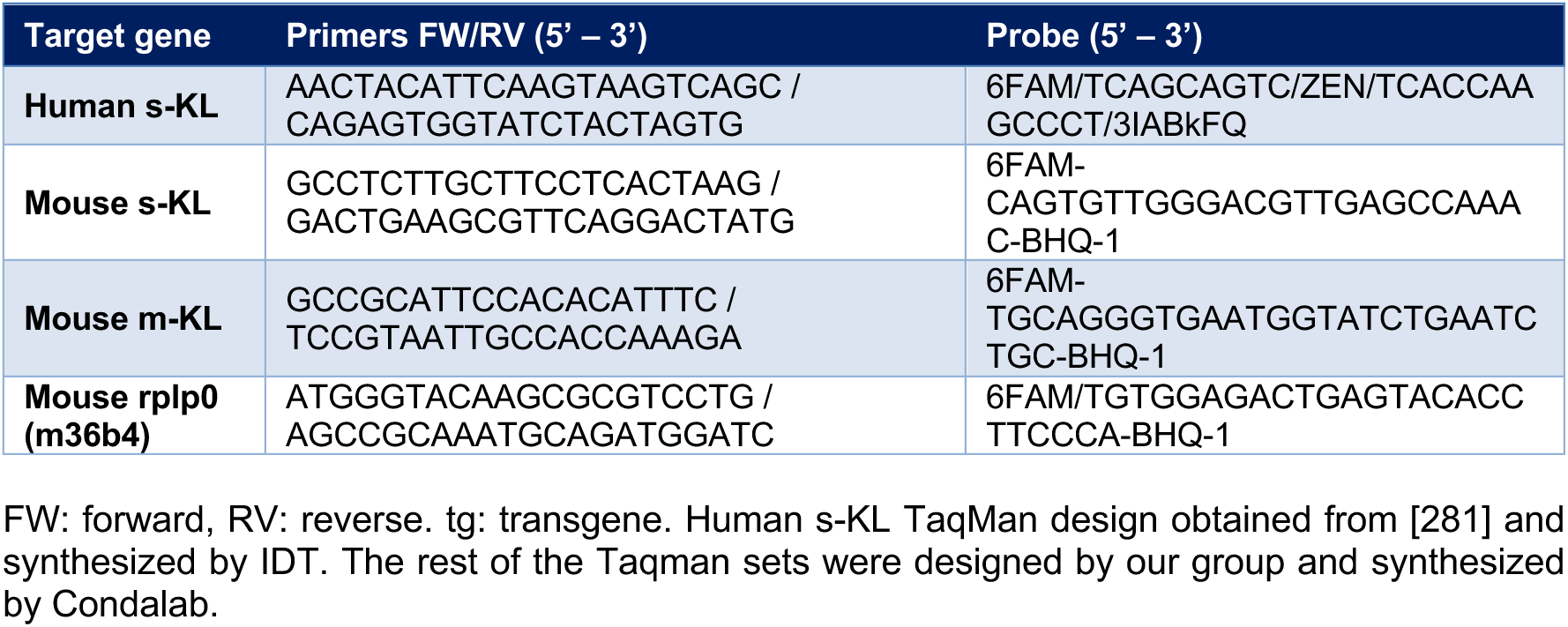
Sequences of custom design Taqman sets.

#### Cerebrospinal fluid (CSF) samples

CSF samples were provided by Dr. Alberto Lleó from the SPIN cohort at the Hospital Sant Pau (Barcelona, Spain) (Alcolea et al. 2019). The group consisted of samples from individuals cognitively unimpaired (CU), Mild cognitive impairment (MCI), or mild dementia (see **¡Error! No se encuentra el origen de la referencia.**).

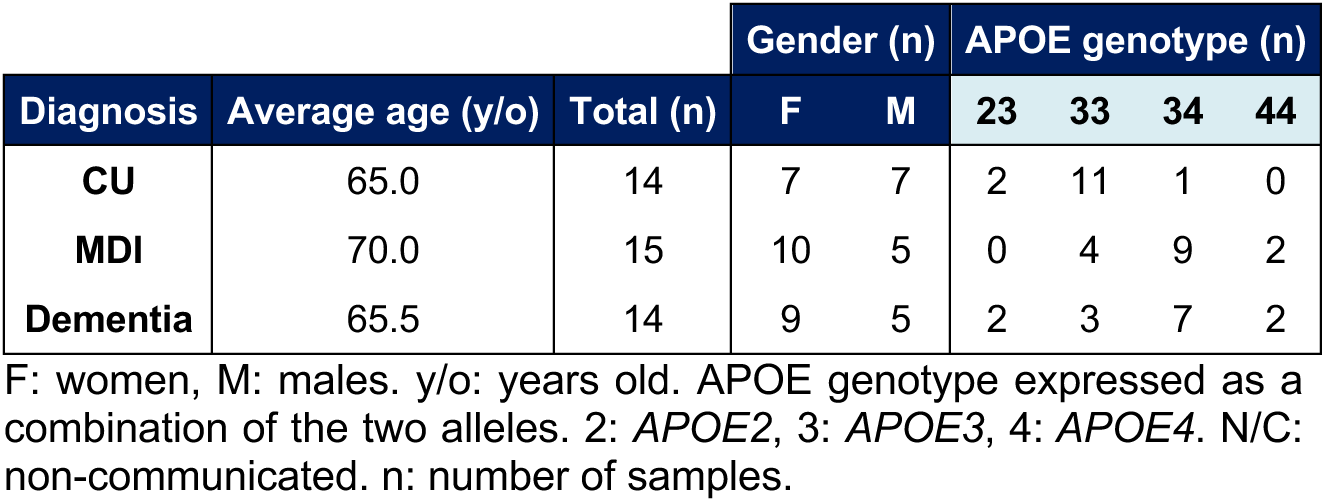

### Mouse, lemur and macaque samples

Brain samples from female and male wild type (WT) mice with genetic background C57BL/6 were extracted at 9 months old. Animals were euthanized by cervical dislocation; the skull was removed, and brain regions were dissected on a cold surface. The frontal cortex, entorhinal cortex, hippocampus and cerebellum were snap-frozen in dry ice and stored at −80°C until RNA extraction. This study was carried out in strict accordance with the current National regulations. The Committee on the Ethic of Animal Experiments of the Universitat Autònoma de Barcelona approved the procedures described (protocol number: CEEAH 4882).

Eight lemurs were used in this study. The animals were born and raised within our breeding facility in Montpellier, France (license approval N°34-05-026-FS). The study was conducted according to the guidelines of the European directive 2010/63 and approved by the Ethics Committee Occitanie Méditerranée CEEA-LR36 (authorization #CEEA-LR-12060). Brain was removed and bisected sagitally, then the left hemisphere was snap frozen and stored at −80°C until processed RT-qPCR studies. The right hemisphere was fixed for 24 hours in 4% PFA and then placed in phosphate buffer. Samples were then embedded in paraffin wax. Brain sections were cut on a sliding microtome at a thickness of 6 μm for histological studies. The primary antibody, mouse monoclonal klotho antibody F-5 (Santa Cruz Biotechnology, ref sc-515939), diluted 1/100 in blocking buffer, was applied overnight at 4°C. Immunological complexes were detected by sequential application of a 1/500 dilution of biotinylated goat anti-mouse immunoglobulin (Vector laboratories, Bur-lingame, CA, USA), and avidin-biotin-peroxidase complex (1/500 dilution; Vectastain ABC kit, Vector laboratories, Burlingame, CA, USA) for 30 minutes each. Staining development was performed with 3,30-diaminobenzidine (DAB kit, Vector laboratories, Burlingame, CA, USA).

A total of three macaques (*Macaca fascicularis*) were used for this study, two 42 months-old females with a body weight of 2.9 and 3.1 Kg and one 36 months-old male with a body weight of 4.2 Kg. Animal handling was conducted at all times following the European Council Directive 2010/63/UE and in full keeping with the Spanish legislation (RD53/2013). The experimental design was approved by the Ethical Committee for Animal Testing of the University of Navarra (ref: 016-17) and by the Department of Animal Welfare of the Government of Navarra (ref: GN016-2017). Anesthesia was induced with an intramuscular injection of ketamine (10 mg/Kg) followed by a terminal overdose of sodium pentobarbital 200 mg/Kg). Animals were transcardially perfused with 1,000 ml of a sodium phosphate buffered solution at neutral pH. Next, the brains were removed from the skull and 4 mm-thick coronal sections covering the whole rostrocaudal extent of the brain were prepared on a monkey brain matrix for cynomolgus monkeys (AST Instruments, MI 48089, USA; Ref. MBM-2500C) on a cold surface and immediately frozen on dry ice. Next, disposable biopsy punches (Kai Europe GmbH, 42697 Solingen, Germany; Ref: BP-40F) were used to collect cylinder-shaped brain samples (4 x 4 mm in size) across different brain areas of interest and stored at −80 °C until RNA extraction was performed. Samples were taken from the superior frontal gyrus (SFG), precentral gyrus (PrG), entorhinal cortex (Ent), hippocampus (Hipp), and cerebellar hemispheres.

### Taqman sets for gene expression analysis

TaqMan sets (composed of a pair of primers and a probe), used to analyze gene expression of s-KL and m-KL in humans and mice were either custom-designed and synthesized by IDT (see Table 4) or inventoried by Invitrogen, Thermo Fisher Scientific (see Table 5). Primers to analyze gene expression of s-KL and m-KL in rhesus monkey were custom-designed and synthesized by Invitrogen.

**Table 5.**
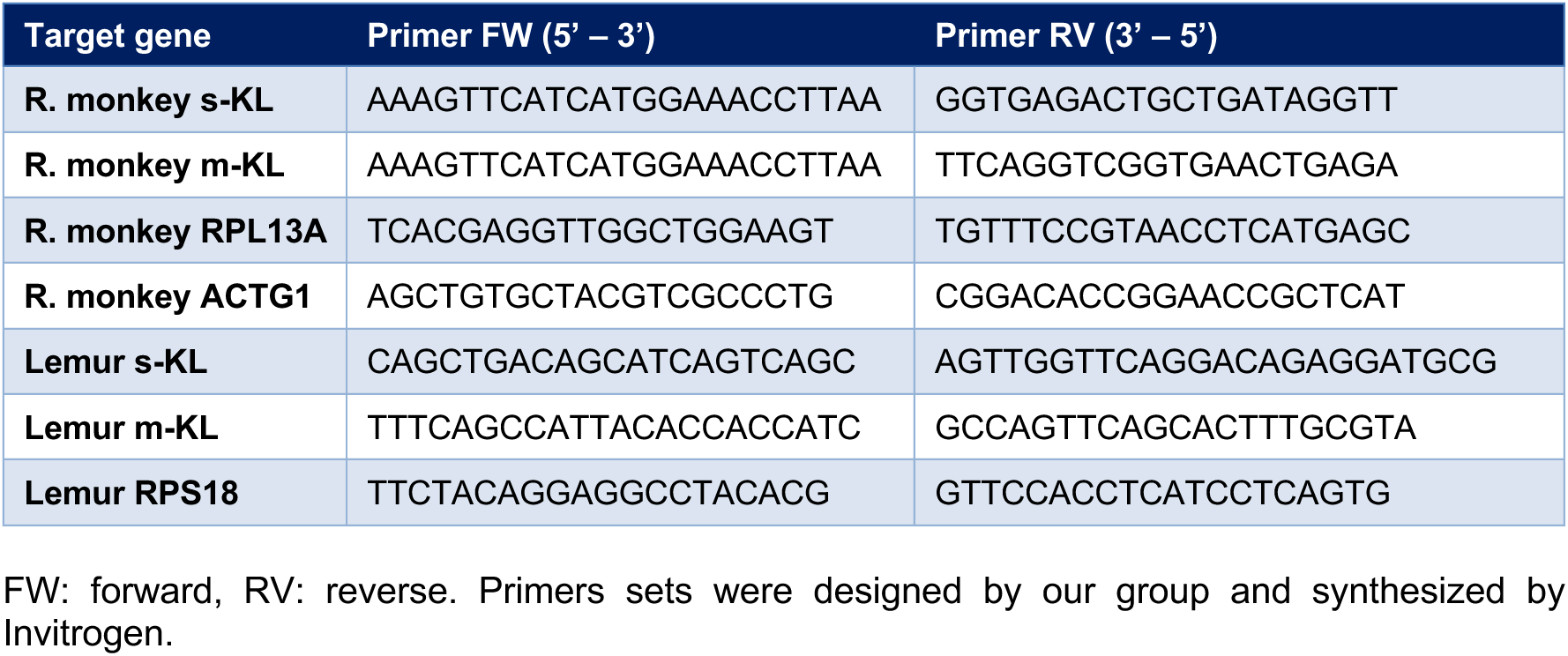
Sequences of custom design primers sets.

**Table 6.** Taqman sets inventoried by Invitrogen (Thermo Fisher Scientific).

| Target gene | Reference |
| --- | --- |
| Human m-KL | Hs00935388_m1 |
| Human UBE2D2 | Hs00366152_m1 |
| Human CYC1 | Hs00357717_m1 |

### RNA isolation and gene expression analysis

To prevent RNA degradation, surfaces, materials, and gloves were cleaned with 70% ethanol and RNAseZap (Ambion), RNAse-free water and solutions were utilized, and samples were maintained at cold temperatures throughout. Human and mouse brain samples were homogenized using a TissueLyser (QIAgen) with grinding beads in the presence of QIAzol Lysis Reagent (Qiagen) for RNA isolation. RNA concentration was measured using the NanoDrop-1000 spectrophotometer (Thermo Scientific) before reverse transcription with SensiFast cDNA synthesis kit (Bioline).

Gene expression was analyzed by real-time quantitative PCR (RT-qPCR) in the CFX384 Thermocycler (BioRad) at the Analysis and Photodocumentation Service of the UAB. Each qPCR reaction consisted of 50 ng of cDNA template, 5 µM Taqman probe, 5 µM primers, SensiFAST Probe No-ROX One-Step Kit (Bioline) and MilliQ water to a final volume of 10 µL in a Hard-Shell Rhin-Wall 384-Weel Skirted PCR (BioRad) well. Two replicates were implemented per reaction as follows: 50°C for 2 min and 40 cycles of 95°C for 10min, 95°C for 15 s and 1 min at 60°C. The average cycle threshold (Cq) of replicates was calculated for each reaction, and samples with replicates having a Cq difference of 1 or higher were discarded. Cq values above 35 were considered as no detectable gene expression and were also discarded. Gene expression quantification relative to two reference genes was calculated using Hellemans’ relative quantification model for normalization with multiple reference genes [283]. UBE2D2 and CYC1 were utilized as reference genes in human samples, and rplp0 for mouse samples, RPS18 for lemur samples and both RPL13A and ACTG1 for R. monkey samples.

The relative expression of s-KL versus m-KL was calculated for each individual using the ratio s-KL/m-KL, considering the difference between the Cq values of the two transcripts and assuming equivalent amplification efficiencies of the TaqMan sets because they all have amplification efficiencies very close to 100%:

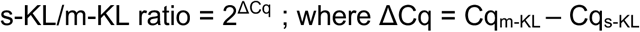

In cases where m-KL was more abundant than s-KL, the inverted ratio (m-KL/s-KL) was calculated to obtain integer values:

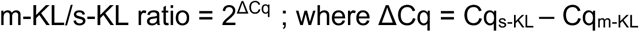

Subsequently, the average s-KL/m-KL (or m-KL/s-KL) ratio was calculated per group.

### Tissue processing for protein quantification

Total protein was extracted from different human brain areas by sample solubilization using lysis RIPA buffer containing 25mM HEPES, 0.2% Igepal, 5mM MgCl2, 1.3 mM EDTA, 1 mM EGTA, 0.1 M PMSF and protease inhibitor cocktails (1:100, P0044; Sigma-Aldrich) at 4°C using a sonicator. Then, samples were centrifuged at 13000 rpm for 5 min at 4°C and the supernatants collected. Total protein concentration was assessed with a commercial Pierce BCA Protein Assay kit (23225; ThermoFisher Scientific) according to manufacturer’s protocol. Protein lysates were aliquot and stored at −80°C until used for analysis.

### Quantification of Klotho by ELISA

Human total soluble KL and secreted-KL levels were quantified using the commercial Human soluble Alpha Klotho ELISA kit (27998, IBL, Minneapolis, USA) and human secreted Klotho ELISA kit (27901, IBL, Minneapolis, USA) respectively, following the manufacturer’s instructions. CSF from human patients was diluted 1:4 in EIA buffer supplied by the kit. For human tissue 100µL of standards and each brain extract (1.0 µg/µL total protein) were added to wells and incubated overnight at 4°C. After washing with wash buffer, the plate was incubated with 100µL of labelled antibody for 60 min at 4°C, followed by TMB solution for 30 min at 4°C. Finally, MAGPIX device with the xPONENT 4.2 software was used to read the plate at 450nm wavelength. Data were expressed as pg/mL of protein. Unfortunately, the ELISA kit for total soluble KL was not functional in brain parenchyma areas.

### Immunoprecipitation and Western Blot analyses

To isolate the secreted Klotho (s-KL) protein from hippocampus, the Pierce® Crosslink Immunoprecipitation Kit (ThermoScientific, ref.26147) was used following the manufacturer’s protocol. Five immunoprecipitation columns were prepared: a negative control column and columns with antibodies BAF1819 (R&D Systems), KM2060 (TransGenic), K113 (Masso et al.(Masso et al. 2015)). Of note, each antibody was used at a concentration of 1 μg/μL in 100 μL, incubated with the agarose resin for 1 hour, and crosslinked using a 2.5 mM DSS solution. Because the efficiency of the K113 antibody was lower than the two commercial BAF1819 and KM2076 antibodies, a fourth condition using 10 times more K113 antibody was also used. Protein samples (1 mg) were pre-cleaned using control agarose columns before incubation with the antibody-bound agarose columns for approximately 24 hours. The immunoprecipitated proteins were eluted and stored for further analysis.

Proteins were resolved on 10% polyacrylamide gels, at 100 V for 20-25 minutes, followed by 1-1.5 hours at 130 V, and transferred to PVDF membranes (GE Healthcare) in a Trans-Blot semi-dry transfer system (Bio-Rad) and run for 45 minutes at 25 V. Membranes were blocked in 5% BSA for 1 hour with constant agitation, washed 3 times and incubated with primary antibodies diluted in 5% BSA-TBST for 1 hour at room temperature or overnight at 4°C. For BAF1819, the antibody was diluted in 5% milk-TBST. After incubation, membranes were washed three times with TBST and incubated with the appropriate secondary antibody for 1 hour. For BAF1819, a third incubation with streptavidin-HRP was performed. Also, in order to allow better detection of faint bands, the membrane incubated with the antibody K113 was also overexposed, previous removal of the 10X-K113 lane. Chemiluminescence detection was carried out using the ETA C ULTRA 2.0 kit (Cynagen) following the manufacturer’s instructions. Protein bands were visualized using the ChemiDoc MP imaging system (Bio-Rad)

**Table 6.** Antibodies and Western Blot Conditions.

| Antibody | Host | Dilution | Reference |
| --- | --- | --- | --- |
| α-Klotho (biotinylated) | Goat | 1:1.000 | RDsystems (BAF1819) |
| α-Klotho KO603 | Rat | 1:500 | TransGenic (KM2076) |
| α-s-KL K113 | Rabbit | 1:10.000 | Masso et al.(Masso et al. 2015) |
| α-Goat Ig HRP (secondary) | Rabbit | 1:10.000 | Dako (P0160) |
| α-rabbit Ig HRP (secondary) | Pig | 1:10.000 | Dako (P0399) |
| α-rat Ig HRP (secondary) | Sheep | 1:10.000 | Thermofisher (PA1-28642) |

### Double immunofluorescence of klotho variants in human tissues

Parallel cryostat slides (12-µm thick) from different human brain areas (hippocampus and amygdala) were obtained using a CM3050s Leica cryostat and processed for double immunohistochemistry against s-KL with either klotho total (KL) or GFAP. Briefly, slides were fixed for 5 min in 4% paraformaldehyde, washed with TBS and inhibited for endogenous peroxidase by incubating samples with a solution of 2% H_2_O_2_ in 70% methanol for 10 min. Then, sections were incubated for 1h in blocking solution containing 10% foetal calf serum in TBS with 1% Triton X-100 (TBS-T) and incubated with rabbit anti-s-KL (1:100, KL182 produced by EZBiolab using the designed immunogenic peptide VSPLTKPSVGLLLPH as antigen) diluted in the blocking solution overnight at 4°C plus one additional hour at RT. After washing three times with TBS-T, sections were incubated with AlexaFluor 555 anti-rabbit antibody (1:500; A21434; ThermoFischer Scientific) diluted in blocking solution for 1h at RT and washed with TBS-T. Then, sections were incubated again for 1h with blocking solution, and then, with either rabbit anti-KL primary antibody-Alexa 488 conjugated (1:100; PA-88303; Thermo Scientific) or with mouse anti-GFAP (1:500, G3893, Sigma) diluted in the blocking solution overnight at 4°C plus one additional hour at RT. KL antibody was conjugated with Alexa 488 using FlexAble CoraLite®488 Antibody Labelling kit (KFA001; Proteintech) following the manufacturer’s instructions. Sections were washed with TBS and then TB; and counterstained with DAPI (1:10,000; D9542; Sigma-Aldrich) diluted in TB for 5 min. Finally, sections were mounted on gelatin-coated slides and coverslipped with Fluoromount-G Mounting Medium (00-4958-02; ThermoFischer Scientific). Negative controls were performed using sections incubated in media lacking the primary antibodies. An additional negative control incubating the slides with s-KL that was previously incubated with the recombinant s-KL protein was used to ensure the specificity of the customer s-KL antibody. Sections were analyzed and photographed with a Zeiss LSM 700 confocal microscope.

### Double immunofluorescence of klotho variants in mouse tissues

Immunofluorescence staining amplified by Tyramide Signal Amplification (TSA) was conducted on 16 µm sagittal cryosections of wild-type (WT) mouse brains using the ThermoFisher kit (ref. B40933). Briefly, brain slices underwent antigen retrieval in a water bath set to 95°C for at least 10 minutes with citrate buffer (10mM citrate, pH 6.0). Following washes with 1x phosphate buffered saline (PBS), hydrogen peroxide solution was applied to inhibit endogenous peroxidase activity for 1 hour. After additional PBS washes, endogenous streptavidin and biotin blockage was performed using the Invitrogen Kit K (ref. E21390). Subsequently, samples were permeabilized with 0.3% Triton X-100 in PBS, followed by blocking with 10% horse serum and 0.05% Tween-20 in PBS (ThermoFisher, ref. 16050122) for 1 hour. Primary antibodies, including anti-s-KL (1:400, K113, in-house (Masso et al. 2015)), anti-KL (1:100, PA5-88303, Invitrogen), and anti-NeuN (1:400, ABN91, Millipore), were diluted in blocking buffer and incubated overnight at 4°C. The next day, slides were washed with 0.3% Triton X-100 in PBS and then incubated with secondary antibodies, biotinylated horse anti-rabbit IgG (1:200, BA-1100, Vector) or goat anti-chicken IgY (1:200, A-11039, ThermoFisher), for 90 minutes.

After additional washes with 0.3% Tween-20 in PBS, slices were incubated with Streptavidin-HRP (Str-HRP) for 1 hour, followed by further washes. For TSA implementation, tyramide working solution was prepared according to the manufacturer’s instructions. After 6 minutes and 30 seconds of reaction, the kit stop reagent was added to the slice for at least 5 minutes. Following washes with 1x PBS, slides were mounted with DAPI Fluoromount-G (00-4959-52, ThermoFisher). All washes were performed three times for at least 5 minutes. Images were captured at 20x magnification using confocal microscopy (Zeiss LSM 700), and colour balance adjustments were made using ImageJ software (NIH).

### Statistical analysis

Statistical analyses and graphical representations were performed using GraphPad Prism software. Differences were assessed by t-test for two means and One-way ANOVA followed by either Tukey or Dunnett’s post hoc tests (as indicated in figure legends) for comparisons involving more than two means. Differences in s-KL versus m-KL expression were analyzed by the One-sample t-test with a null hypothesis equal to 1. Pearson correlation coefficient (r) was calculated to assess linear correlation between two continuous variables. Null hypotheses were rejected at or below a p-value of 0.05.

## Acknowledgements

We are thankful to the personnel of the animal facility of the lemur colony at RAM-CECEMA, Montpellier. This work was funded by MICINN/AEI/10.13039/501100011033 (PID2022-142624OB-100) to MC; and (PID2020-116735RB-I00 and PID2023-148834OB-100) to AB; by Instituto de Salud Carlos III and NexGenerationEU (RICORS, RD24/0014/0039) to MC and AB; and Generalitat de Catalunya (2021SGR-00529) to AB and MC. RB was a recipient of a fellowship by the Gobierno de España (FPU16/03137).

## Author Constributions

R.B. and M.C. conceptualized and designed the study. R.B., J.E., N.MF. and G.D. performed the in vivo experiments; R.B., R.GY., J.dR., J.RS., N.MF., G.D. analyzed the samples; R.B. R.GY., J.RS., N.MF., B.A. and A. B. interpreted the experimental data; A.L., E.C., A.N., I.F. and JL.L. contributed, genotyped and/or phenotyped, and characterized human and non-human primate samples. R.B. and M.C. wrote the manuscript with the help of the other authors. M.C. secured primary funding, supervised and coordinated the project.

